# Spatial hearing and temporal processing ability of the Mongolian gerbil (*Meriones unguiculatus*) measured using prepulse inhibition of acoustic startle

**DOI:** 10.1101/2025.03.12.642882

**Authors:** Matthew D. Sergison, Shani Poleg, Nathaniel T. Greene, Achim Klug, Daniel J. Tollin

**Affiliations:** Neuroscience Graduate Program, University of Colorado School of Medicine, Aurora, CO 80045; Department of Physiology and Biophysics, University of Colorado School of Medicine, Aurora, CO 80045; Department of Otolaryngology, University of Colorado School of Medicine, Aurora, CO 80045

## Abstract

The Mongolian gerbil is a common model organism for studying the neural and behavioral mechanisms of binaural and spatial hearing, largely because of its ability to hear lower frequencies than other rodents and thus utilize both interaural time and level difference cues for sound localization. Prior spatial hearing studies in gerbils have relied on operant conditioning paradigms, requiring large amounts of time-consuming training and testing on multiple different tasks needed to make a comprehensive assessment of spatial hearing ability (including temporal processing, spatial acuity and spatial unmasking). This limits the ability of researchers to thoroughly assess behavioral performance in individual animals. In this study, we used the reflexive prepulse inhibition of the acoustic startle reflex (PPI) to extensively assess spatial hearing and temporal processing abilities in individual gerbils of both sexes. Results show that gerbils inhibit a startle response to a brief loud sound based on prepulse acoustical cues consisting of a 1) temporal gap in ongoing sounds, 2) change in sound source location, and 3) target sound in the presence of a masker. In each test, the magnitude of the suppression of startle increased monotonically as a function of the magnitude of the acoustical prepulse, not unlike a psychometric function, from which threshold performance could be measured. Thresholds in the gerbils in each task measured using PPI matched those acquired using operant conditioning methods.

## 1. Introduction

The Mongolian gerbil (*Meriones unguiculatus*) has become a commonly used animal model in auditory neuroscience research due to its extended low-frequency hearing ability as compared to other commonly used rodents. Gerbils have a low-frequency hearing limit reaching 0.1 kHz, compared to 1.5 kHz in mice and 0.5 kHz in rats (Ryan, 1976) (Turner et al., 2005). This makes them a potentially more translational model for human auditory processing, which mostly occurs in lower frequencies (20Hz-20kHz) (Turner et al., 2005). Gerbils also allow for high throughput of experiments, as well as experiments that require larger numbers of subjects as compared to other common auditory animal models, such as cats or non-human primates.

The gerbil’s low frequency hearing makes them an ideal model organism for studying the neural and behavioral mechanisms of binaural hearing and sound source localization. In mammals, low frequency sounds, namely lower than 2kHz in gerbils (Tolnai et al., 2017) (R. S. Heffner & Heffner, 1988b), are localized on the basis of interaural time differences (ITDs) in the fine structure of ongoing sound waves, and ITDs are initially encoded by neurons comprising the medial superior olive (MSO) (Yin & Chan, 1990) (Goldberg & Brown, 1969) (Pecka et al., 2010). Conversely, high frequency sounds, or higher than 2kHz in gerbils (Tolnai et al., 2017) (R. S. Heffner & Heffner, 1988b), are localized by interaural level differences (ILDs), and ILDs are initially encoded by neurons comprising the lateral superior olive (LSO) (Grothe et al., 2010) (Tollin, 2003) (Yin et al., 2019). Neurons of the LSO also encode ITDs in transient sounds (e.g., snapping twigs, etc.) and the low-frequency envelopes of high-frequency sounds (Joris & Yin, 1995) (see (Owrutsky et al., 2021) for review of the different types of ITDs and their encoding by the LSO).

In order to use the gerbil as a model system for binaural and spatial hearing studies, a behavioral paradigm that shows their capability to not only localize sound sources but also to use binaural cues for other behavioral tasks is necessary. Several prior studies have used traditional operant conditioning paradigms to measure the gerbil’s sound localization ability (Carney et al., 2011) (Lingner et al., 2012) (Maier & Klump, 2006) (Tolnai et al., 2017) (Lesica et al., 2010) (Maier et al., 2008) (R. S. Heffner & Heffner, 1988b). For the most part, these have been studies of spatial acuity, i.e. the smallest change in sound source location that the animal can just detect, often referred to as the minimum audible angle (Mills, 1958). For broadband noise sources, these studies have shown that gerbils have spatial acuity ranging from ∼25-38°.

However, operant conditioning experiments require long periods of training prior to collecting data, followed by many weeks of repeated experimentation to obtain reliable data. Moreover, training on one task, such as spatial acuity, is not generalizable to performance on a different binaural task, for example detecting a signal in spatially separated noise, or tasks of temporal processing, such as gap detection. Because of this, traditional methods such as operant conditioning are not likely to be as useful for in-depth studies of conditions where binaural hearing is impaired, such as aging (Laumen et al., 2016), noise exposure (Benson et al., 2025), neurodegenerative diseases (e.g. Alzheimer’s (Kurylo et al., 1993)), or developmental disorders (e.g. Fragile X (McCullagh et al., 2020)). An inability to test a multitude of tasks in the same subjects is significant, as human studies have shown that several kinds of listening difficulties related to the conditions listed above arise not only with binaural and spatial hearing, but also with mechanisms of general temporal processing. To circumvent this, we describe prepulse inhibition (PPI) of the acoustic startle as a behavior metric for high-throughput study of binaural hearing and sound localization. This method requires no training, opening the possibility to test individual animals on a number of different listening tasks.

PPI methods have been used previously in the gerbil. When presented with a loud stimulus, gerbils will reflexively startle (Gaese et al., 2009); however, if presented with a secondary stimulus, or pre-pulse cue that is detectable by the animal just prior to the presentation of the startle stimulus, gerbils will reflexively inhibit their startle (Gaese et al., 2009) with the magnitude of the startle suppression scaling proportionally to the detectability of the pre-pulse cue. The neural circuitry of both the startle and the PPI pathway includes auditory nuclei in the brainstem and midbrain that are directly involved in binaural processing, indicating that sound localization cues could likely be used as the pre-pulse cue to induce PPI (Koch, 1999).

The extent to which gerbils can utilize PPI as a metric for spatial hearing is unclear. Here, we show that gerbils exhibit PPI to acoustic startle in response to a number of auditory pre-pulse cues, including binaurally processed stimuli, and that PPI can be used as a behavioral metric to indicate binaural processing abilities with performance generally matching that obtained via operant conditioning paradigms.

## 2. Methods

### 2.1 Animal subjects

10 Mongolian Gerbils (4 male, 6 female) between the ages of 2-10 months old were used. Animals come from our in-house breeding program. The 2-10 month range of gerbils is commonly used as a young adult timepoint, and it represents an ideal age for testing because the auditory system is fully developed, but not yet undergoing age-related hearing loss or inner ear synaptic degeneration (Steenken et al., 2021). Gerbils were either individually, pair or group housed, and given food and water *ad-libitum*. All gerbils were weighed before each testing session to assess stress levels, and animals were monitored daily by lab personnel or veterinary staff: all animals used were of normal weight and were otherwise healthy. Experimental procedures were approved by the University of Colorado Anschutz Medical Campus Animal Care and Use Committees (IACUC) and followed all the National Institutes of Health Office of Laboratory Animal Welfare (OLAW) guidelines.

### 2.2 Apparatus

Experimental methods follow those detailed in (Greene et al., 2018). All experiments were conducted in a double-walled, sound-attenuating chamber (interior dimensions: ∼3 x 3 x 3 m; IAC, Bronx, NY) lined with echo-attenuating acoustical foam. Animals were placed in an acoustically transparent custom-made cage constructed of wire mesh mounted to a flexible polycarbonate platform and raised to speaker height by four polycarbonate (PVC) posts (Fig. 1A). The dimensions of the mesh wire cage were tailored to allow each animal room to move for a startle response, but not enough for the animal to significantly turn its head or to turn around in the cage. The head of the animals was thus consistently oriented towards the midline speaker, or at the speaker positioned 45 degrees off-midline during hemifield tasks (Fig. 7A, see results for more information). All animals were tested in the dark, and were visually monitored using a closed-circuit infrared (IR) camera to ensure the correct orientation was present at all times.

**Fig. 1.**
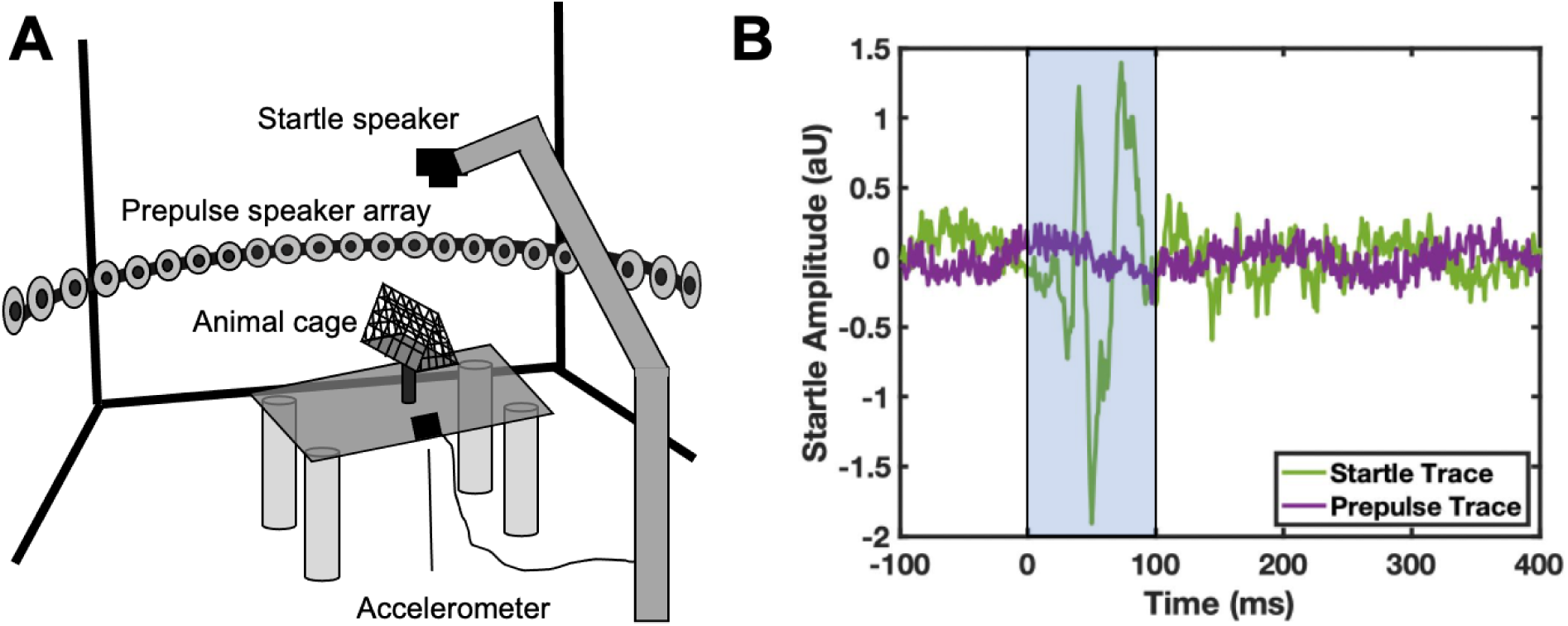
Diagram of PPI experimental setup and example startle traces. **A:** Diagram of our experimental setup for measuring PPI in gerbils. Gerbils sit in the animal holding cage, which is attached to a platform and accelerometer to measure startle amplitude. Prepulse cues are presented from the 180-degree speaker array surrounding the animal. Startle stimuli are presented from the startle speaker located above the animal. **B:** Example traces of startle responses from our experimental setup. Green trace is a control startle condition with no prepulse cue. Purple is from a startle condition with a prepulse cue. Traces are quantified into startle amplitudes by calculating the RMS within the startle window (shaded by blue box).

Sounds that constitute the ‘prepulse’ for the prepulse inhibition of acoustic startle paradigm were presented from one of 25 calibrated loudspeakers (Morel MDT-20) spaced along a 1-m radius semicircular boom at 7.5^°^ increments, from 90° (right) to 90° (left)(see (Greene et al., 2014) for detailed description of the speaker setup). The speaker boom was oriented horizontally (i.e. 0^°^elevation) for all experiments. Startle-eliciting stimuli were presented from a Faital Pro HF102 compression driver mounted ∼25 cm above the animal and amplified with a Yamaha M-40 power amplifier. The startle response of the animal was captured using a cage-mounted accelerometer (Analog Devices ADXL335)(Fig. 1B).

Three Tucker-Davis Technologies (TDT) RP2.1 Real-time Processors controlled by custom written MATLAB (MathWorks) scripts generated the stimuli and recorded the startle response. The startle-eliciting stimulus (SES) was a single 50-millisecond (ms) duration broad-band noise burst (rectangular-gated, 50 kHz bandwidth) dynamically generated by the first RP2.1, and presented at 117 dB SPL, except when testing startle response of gerbils to different startle speaker levels (Fig 2). Carrier stimuli (CS) consisted of broad-band noise dynamically generated by the second RP2.1, and presented continuously (except when otherwise noted) during testing. In some experiments the broad-band noise CS was low- or high-pass filtered with a 100th order FIR filter that was designed in MATLAB and implemented in the second RP2.1. The CS was presented from one speaker at a time, controlled by switching the output from channel 1 to 2 on the third RP2.1 (which received the CS as an input). The speakers corresponding to these outputs were dynamically set by two sets of TDT PM2Relay power multiplexers, controlled by the first and third RP2.1. The motion of the polycarbonate plate resulting from the startle response was transduced by the accelerometer, amplified 55 dB by a TDT MA2 amplifier, and the voltage output sampled at 1kHz by the first RP2.1 (e.g., Fig 1B). The startle response amplitude was calculated as the RMS of the accelerometer output in the first 100-ms period after the delivery of the SES. The apparatus as well as the prepulse and startle stimuli presentation methods were virtually identical to those used in previously run experiments by our laboratory using guinea pigs (Greene et al., 2018) (Benson et al., 2025) and mice (McCullagh et al., 2020). These methods are an adaptation of those described by (Allen & Ison, 2010).

**Fig. 2.**
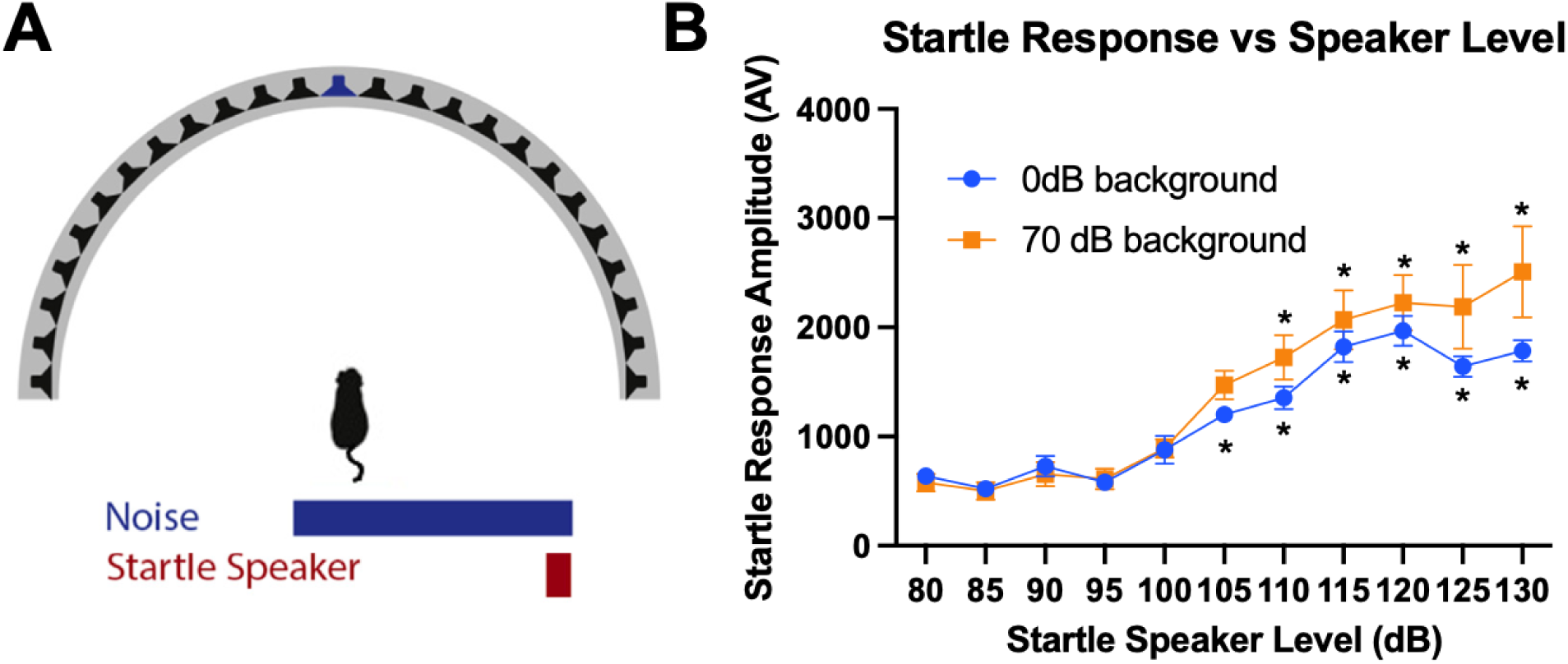
Gerbils elicit a larger startle reaction in response to louder startle stimuli. **A:** Diagram of experimental protocol. Gerbils were held at midline and startle stimuli were presented between 80 and 130 dB SPL in 5 dB increments, either in the presence of 70 dB background noise or in quiet. **B:** Startle amplitudes increased in response to louder startle stimuli presented in both quiet (blue) and 70 dB background noise (orange). Startle amplitudes were significantly increased at 105 dB in quiet and at 110 dB in 70 dB background noise. Asterisks indicate a significantly higher startle amplitude than baseline (80 dB) within each category as determined by a one-way ANOVAs (*p<0.05).

### 2.3 Prepulse inhibition of acoustic startle

Conditions were presented 11 times in all experiments, with the last ten conditions being used to calculate PPI value. The first trial for each experiment was removed to allow for animals to be conditioned to the startle and prepulse stimuli. Animals were limited to <60 mins of testing each day to avoid habituation to the startle stimulus (Benson et al., 2025). In experiments requiring greater than 1 hour of testing, testing was spread out over multiple days, and gerbils were given at least one recovery day in between testing days to further avoid habituation to the startle stimulus. Experimental and control trials within each testing day were presented pseudo randomly in a block-wise fashion (repeating all conditions of a given repetition before moving to the next repetition). Stimulus presentation and response measurement was computer controlled. Each session started with a two-minute acclimation period with no presentation of the startle stimuli. The inter-trial interval (ITI) averaged 20s, which varied randomly between 15 and 25s at an interval of 1s.

For each prepulse task, startle amplitudes from each repetition were averaged for every condition. In tasks where multiple days of testing were performed, measurements were averaged over all days for each condition. Startle amplitude from control conditions (no prepulse cue presented) were averaged together as well. The PPI value for each condition was then calculated as 1-(Startle in Prepulse conditions/Startle in Control Condition). Since PPI value reflects a reduction of startle amplitude, a value closer to 1 indicates higher suppression of the startle response, and therefore better detection of the prepulse stimulus, while values closer to 0 indicate lower suppression of the startle response, and poorer detection of the prepulse stimulus. To quantify this, post hoc tests (after a significant ANOVA test) on PPI values for a condition that is statistically larger than 0 would indicate significant detection of the prepulse stimulus in that condition, while one that is not statistically different from 0 would indicate no detection of the prepulse stimulus. From this type of analysis, we are able to determine the smallest magnitude of the prepulse stimulus that the animals were able to detect, not unlike determining the threshold discrimination from a psychometric function measured using operant conditioning methods.

### 2.4 Startle amplitude response

Before running the animals on any of the prepulse stimulus conditions, the ability of the animals to startle in response to the brief startle stimulus was quantified. Gerbils were held in the apparatus described above with their heads facing the midline (0°) speaker (Fig. 2A). For this test, the decibel level of the startle stimulus varied pseudorandomly from trial to trial between 80 and 130 dB in 5 dB level steps. Each day of testing lasted approximately 45mins. The startle amplitude was measured in response to the different dB levels of the startle stimulus – there was no prepulse stimulus. The startle amplitude response was measured across two days, once in silence, and once in the presence of 70 dB background broadband noise from a speaker above the animal.

### 2.5 Prepulse inhibition of acoustic startle tasks

#### 2.5.1 Gap detection

We used a gap detection task to test 1) the ability of gerbils to utilize a well-studied prepulse cue as a metric of detection and 2) to test the temporal processing abilities of gerbils. Each day of testing lasted approximately 30 mins. In addition to the 7 experimental conditions, 2 control conditions with no prepulse stimulus were intermixed in the testing. In the gap detection tasks, and in all following prepulse tasks, startle stimulus SPL was set to 117 dB SPL ensuring a robust, reliable and repeatable startle amplitude from all of the animals tested. A 70 dB SPL broadband noise was presented from the speaker array. The prepulse cue in this task was a temporal gap of silence presented within the ongoing broadband noise. In the first day of gap detection testing, the gap was a fixed 20-ms, and the timing of the gap presentation varied before the onset of the startle stimulus at 5, 10, 20, 40, 80 or 160-ms (Fig. 3A). A 20-ms gap was chosen here because it has been shown previously to be reliably detected by gerbils (Hamann et al., 2004). The second day of testing was identical to the first day, but with a 50-ms gap in the broadband noise instead of a 20-ms gap (Fig. 3A). The third day used a gap of variable length of either 1, 3, 5, 10, 15 or 20-ms, which was constantly presented 80-ms prior to startle stimulus presentation (Fig. 4A).

**Fig. 3.**
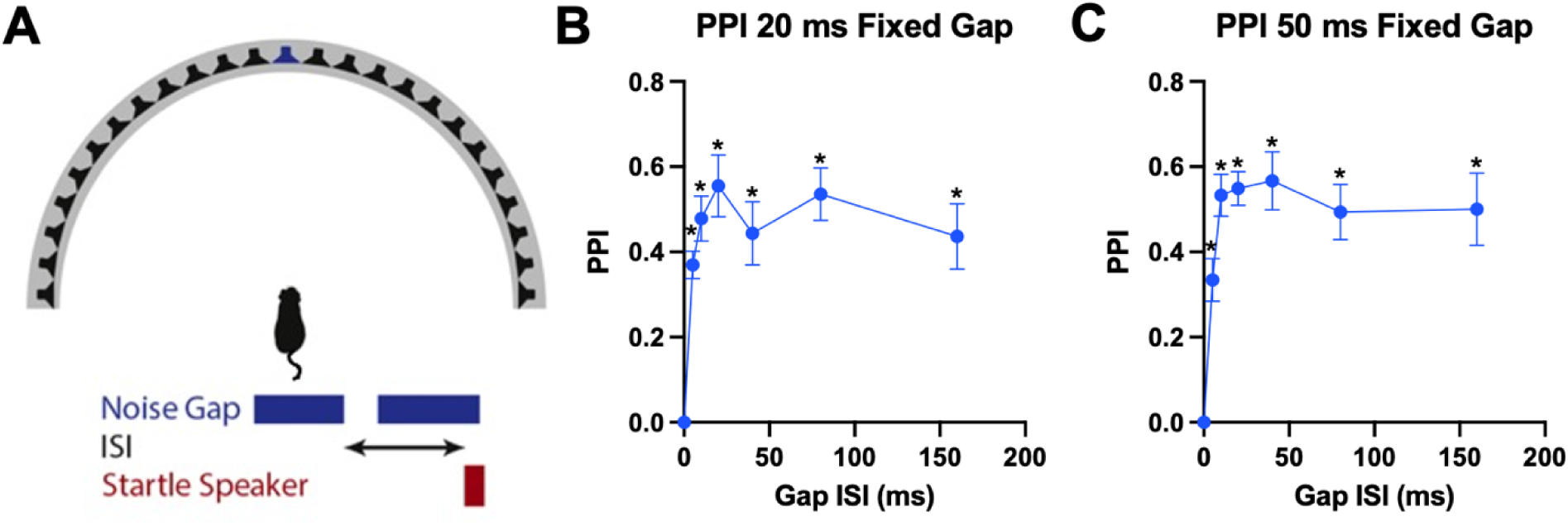
Gerbils can utilize PPI in response to a fixed length gap in an ongoing broadband noise. **A:** Diagram of the fixed gap detection experiment setup. Gerbils were presented a gap of either 20 ms or 50 ms, which varied in time of presentation prior to startle stimulus (inter-stimulus interval, or ISI). **B:** PPI in response to 20 ms gap at different ISIs (5, 10, 20, 40, 80 and 160 ms). Asterisks indicate significant PPI above zero as determined by a one-way ANOVA (*p<0.05). Gaps were detected in all ISIs between prepulse and startle stimuli. **C:** PPI in response to 50 ms gap at different ISIs (5, 10, 20, 40, 80 and 160 ms). Asterisks indicate significant PPI above zero as determined by a one-way ANOVA (*p<0.05). Gaps were detected in ISIs between prepulse and startle stimuli ≥ 10 ms.

**Fig. 4.**
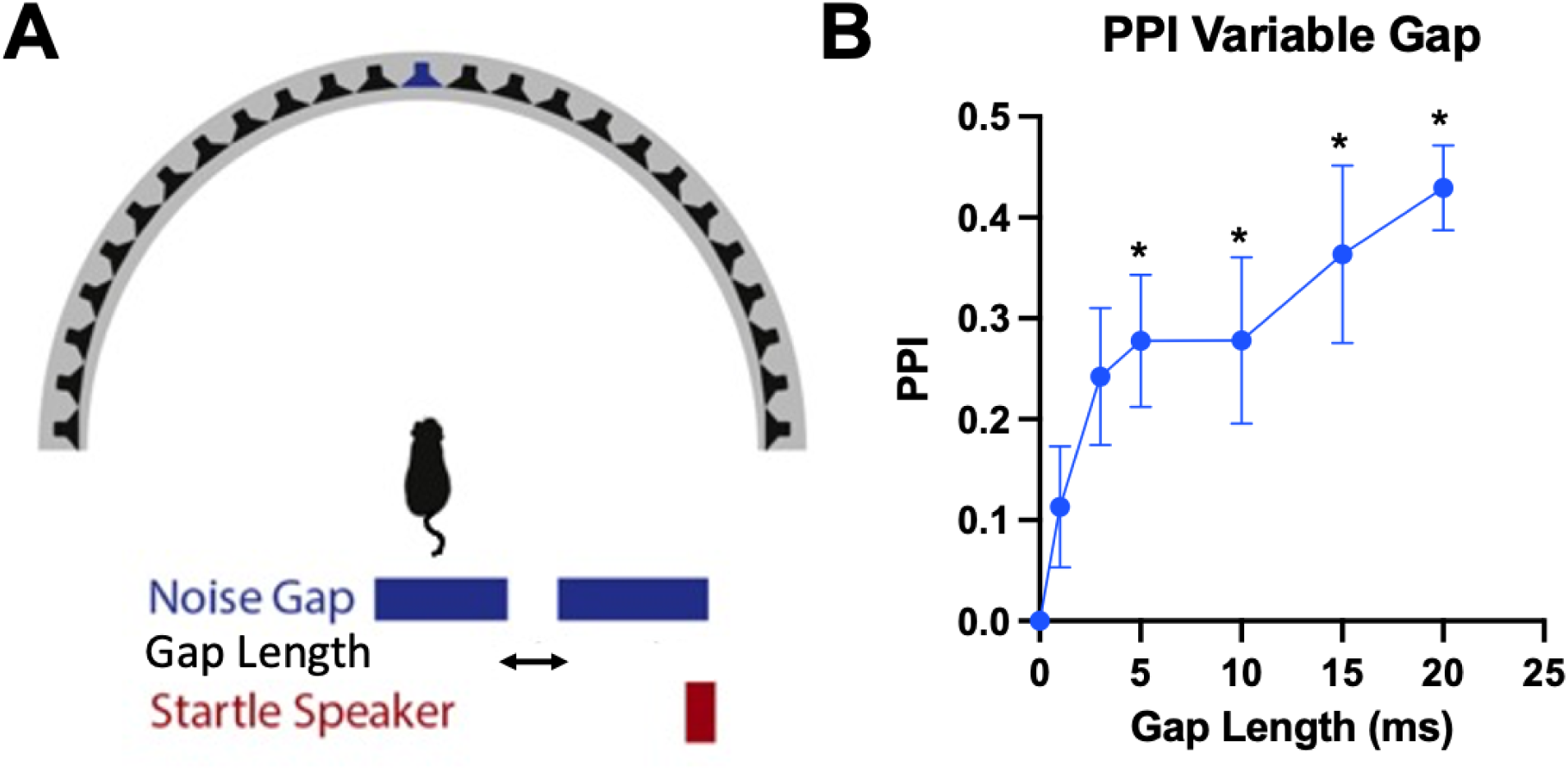
Gerbils can utilize PPI in response to a variable length gap in an ongoing broadband noise. **A:** Diagram of the variable gap detection experiment setup. Gerbils were presented a gap at varying lengths, which was consistently presented 80 ms prior to startle stimulus. **B:** PPI in response to gaps of different lengths (1, 3, 5, 10, 15 or 20ms). Asterisks indicate significant PPI above zero as determined by a one-way ANOVA (*p<0.05). Gaps ≥ 5 ms in length could be detected by gerbils.

#### 2.5.2 Minimum audible angle

We used the so-called speaker swap (Allen & Ison, 2010) paradigm to assess the smallest change in the spatial location of sound sources that the animals could just detect, or the minimum audible angle (Fig. 5A). Each test session lasted <40 mins. Animals were held in the wire mesh cage with their heads oriented towards the 0° midline speaker in the apparatus described above. The prepulse stimulus in the spatial acuity experiments was a change in 70 dB SPL noise sound source location of the sound between two speakers with complementary angles relative to the midline. For example, the stimulus would initially be presented at the speaker at +45° and then swapped to the speaker at -45°. The startle stimulus was presented 80-ms after the swap in speaker location (Fig. 3B, 3C). The stimulus would then be presented from a randomly selected different speaker location for the next trial. The prepulse stimuli consisted of either 1) broadband noise or broadband noise filtered by a 2) low-pass (0.5 kHz) or 3) high-pass (4kHz) filter (Fig. 6A). The low-pass and high-pass conditions test the ability of the gerbils to use low-frequency ongoing ITDs or high frequency ILDs for sound localization, respectively (Greene et al., 2014), (Greene et al., 2018). The highest frequency at which gerbils are able to utilize low-frequency ongoing ITDs for sound localization and begin using high-frequency ILDs is ∼2 kHz (Tolnai et al., 2017) (R. S. Heffner & Heffner, 1988b), so a 0.5 kHz low pass ensures that gerbils must rely almost completely on the use low-frequency ongoing ITDs cues; moreover, measured ILDs in gerbil for frequencies < 2 kHz are negligible (< 3 dB) at any angle along the horizontal plane due to the small size of the head (Maki & Furukawa, 2005). On the other hand, a 4 kHz high pass noise ensures gerbils cannot use fine structure ITDs will have to rely on the use of high frequency ILD cues, which approach ∼30 dB at the largest angles used here (Maki & Furukawa, 2005). In the hemifield experiments, gerbils were tested to see if they could discriminate sound source location within a hemifield, as opposed to sounds that swapped source location across the midline. Human listeners can discriminate changes in sound source location not only across the midline, such as the tasks described above, but also if the sources are located within a hemifield on one side of the head (Mills, 1958); our hemifield speaker swap test was meant to assess if gerbils are able to do the same. Here the same apparatus and conditions as described above were used except the cage was rotated so that the head of the animal was oriented towards the speaker located at either the speaker located at -45° or +45° degrees in the right or left hemifield (Fig. 7A). In all experiments, control conditions, where startle was presented but with no speaker swap prepulse (one control condition for each starting speaker tested), were intermixed with experimental trials in each day of testing. We hypothesize that speaker location swaps that subtended larger angles would be easier to detect, and thus suppress the startle response more robustly resulting in larger PPI values approaching 1.0, compared to more narrow swap angles that would be more difficult to detect, thus producing less startle suppression and smaller PPI values approaching 0.0.

**Fig. 5.**
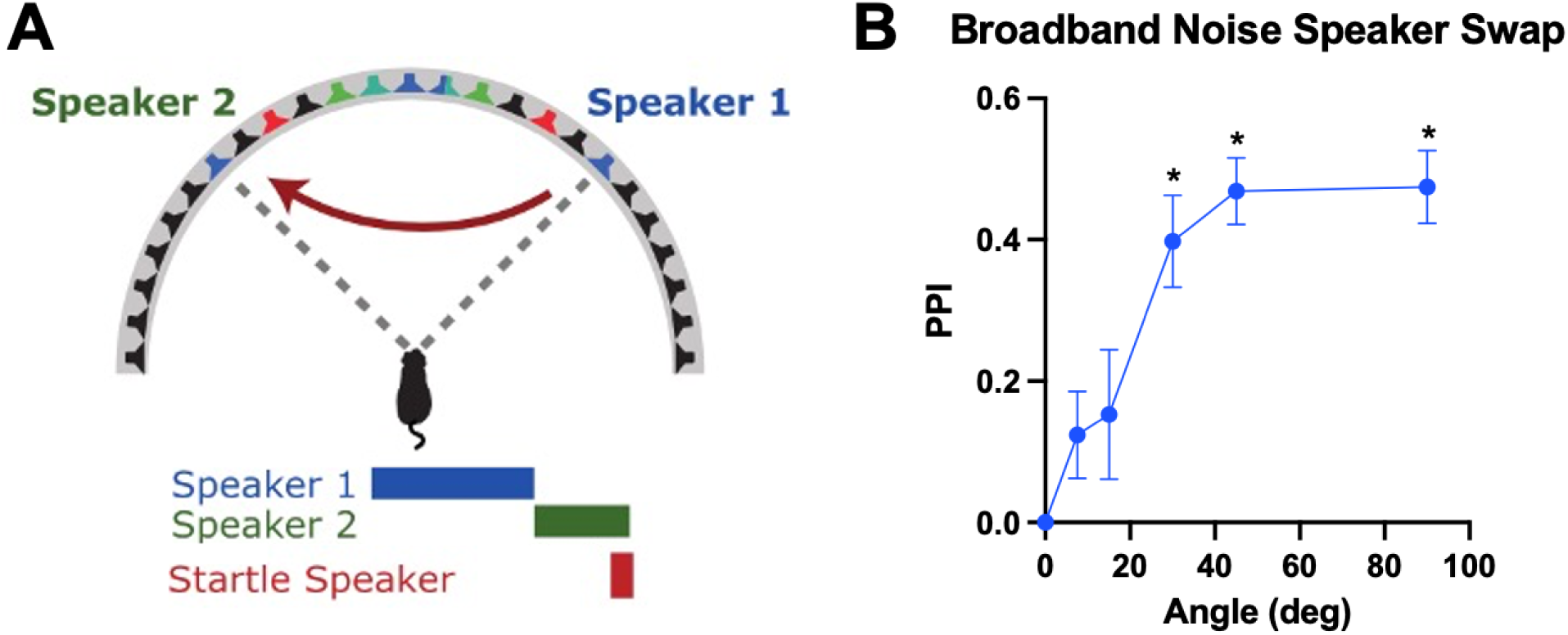
Gerbils can utilize PPI in response to a change in the spatial location of broadband noise. **A:** Diagram of the broadband-noise speaker swap experiment setup. Gerbils were held at midline, while broadband noise was presented at one speaker, then swapped to a corresponding speaker across the gerbil midline 80 ms prior to startle stimulus. The angle of the speaker swap was varied. **B:** PPI in response to different speaker swap angles (7.5, 15, 30, 45 or 90°). Asterisks indicate significant PPI above zero as determined by a one-way ANOVA (*p<0.05). Speaker swap angles ≥ 30° could be detected by gerbils.

**Fig. 6.**
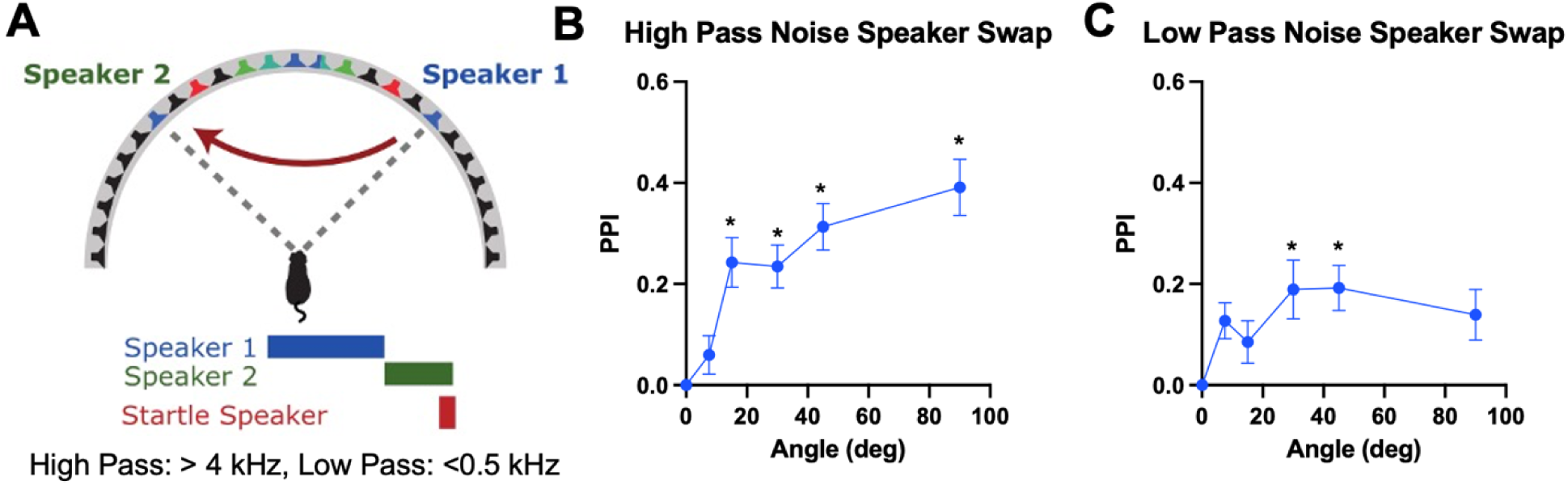
Gerbils can utilize PPI in response to changes in the spatial location of high- and low-pass noise. **A:** Diagram of the speaker swap experiment setup. Gerbils were held at midline, while either noise high-passed at 4kHz or noise low-passed at 0.5 kHz, depending on the experiment day, was presented at one speaker, then swapped to a corresponding speaker across the midline 80 ms prior to startle stimulus. The angle of the speaker swap was varied. **B:** PPI in response to different high pass noise speaker swap angles (7.5, 15, 30, 45 or 90°). Asterisks indicate significant PPI above zero as determined by a one-way ANOVA (*p<0.05). Speaker swap angles ≥ 15° could be detected by gerbils. **C:** PPI in response to different low pass noise speaker swap angles (7.5, 15, 30, 45 or 90°). Asterisks indicate significant PPI above zero as determined by a one-way ANOVA (*p<0.05). Speaker swaps of 30° could be detected by gerbils.

#### 2.5.3 Spatial release from masking

We used a spatial release from masking task to assess the ability of gerbils to detect a stimulus in the presence of a background noise masker (Fig. 8A). The prepulse cue was a series of 8 12.5-ms duration broadband chirps presented from the midline 0° speaker. The chirps were presented at three different levels, 56, 50 or 44 dB SPL. The masking noise was 70 dB SPL. In addition to varying the level of broadband chirp signals, the spatial location of the masker was also varied relative to the midline of either ±7.5, ±22.5 or ±37.5. Each day of testing was <60 mins, with 6 trials per condition being tested and 5 trials being recorded per day. Broadband chirps were always presented 80-ms before startle speaker presentation. After data was collected, we combined the trials of corresponding angles across the midline, or the same angle of separation between masker and signal to the left or right of the animal, for further analysis. Similar to the tasks above, larger values of PPI would indicate 1) better detection of the signal in the noise and therefore 2) better separation and segregation of the signal from the masker.

### 2.6 Statistical analysis

Statistical Analysis was performed using GraphPad Prism V10. We assumed the data to be normally distributed with the 10 animals, but this was not formally tested; our sample size is similar to previous publications (Maier & Klump, 2006) (Carney et al., 2011) (Lesica et al., 2010). Regardless, the *F*-test has been shown to be robust, in terms of Type I error, to slight, moderate, and severe departures from normality, with various sample sizes (equal or unequal sample size) and with same or different shapes in the groups (Blanca et al., 2017). For changes in startle amplitude, we performed one-way ANOVAs on both 0 dB and 70 dB background noise conditions to assess the change in startle amplitude compared to startle stimuli level. An additional two-way ANOVA was performed to assess the difference between startle amplitudes in quiet and in the presence of a 70 dB background noise. For gap detection and speaker swap tasks, one-way ANOVAs were used to compare each experimental condition of different gap ISI, gap duration or speaker swap angle against a control condition of value zero PPI. Detection of the prepulse cue by the animals was determined if the ANOVA showed a significant difference between the conditional PPI value and the zero PPI control. Additionally, one-way ANOVAs were run on each task to look for significant changes between PPI values in different conditions. For spatial release from masking, a two-way ANOVA was performed on the data. This allowed us to look at joint effects of target-to-masker angle and target signal attenuation level on PPI value. Additional one-way ANOVAs were performed for each signal level to assess the spatial differences of each condition. For all ANOVAs, post hoc Bonferroni corrected t-tests were performed to examine which conditions had significant pairwise differences in each experiment.

## 3. Results

### 3.1 Startle response amplitudes increase with increasing startle stimulus level

We first tested the startle threshold in both quiet (0 dB background noise) and in background noise (70 dB broadband noise presented from the midline 0° speaker) (Fig. 2). This task was used to confirm that the animals produced a startle and to determine the startle stimulus SPL level that elicited a robust, reliable and reproducible startle to be used for the subsequent PPI tasks.

Fig. 2B shows the startle amplitude (in arbitrary units) as a function of startle stimulus dB SPL in both the 0 dB and 70 dB background conditions. Overall, startle amplitude increased monotonically with SPL for both conditions above baseline starting at around 105-115 dB SPL. In the 0 dB background condition, the one-way ANOVA indicated that there was a significant effect of startle SPL on startle amplitude [F(10,99) = 30.34, p < 0.0001]. Post hoc tests revealed that startle amplitude was significantly larger than the startle to baseline level (80 dB SPL) in response to all levels ≥ 105 dB (all p’s < 0.0056). Similarly, in the 70 dB SPL background noise condition there was also a significant effect of startle SPL on startle amplitude [F(10,99) = 11.85, p < 0.0001] and post hoc tests revealed a significantly larger startle amplitude compared to 80 dB baseline in response to all levels ≥ 110 dB (all p’s < 0.0206). A two-way ANOVA also showed there was a significant effect of background noise on startle amplitude where gerbils startled more to the same startle stimulus dB level presented within 70 dB background noise compared to no added background noise [F(1,10) = 8.034, p = 0.0051]. Specifically, the startle amplitude was larger in background noise than in quiet at the loudest startle conditions tested, 125 dB (p = 0.0276) and 130 dB (p = 0.0037).

For the prepulse inhibition experiments, we selected the startle stimulus that reliably and reproducibly evoked a statically significant startle above baseline, but not so loud as to potentially induce hearing loss over the course of testing. Based on the results shown above, we thus used 117 dB as the level for the startle stimulus in all remaining experiments as this was the level after which startle amplitude plateaued with even louder startle stimulus SPL.

### 3.2 Gap detection

#### 3.2.1 Fixed gap durations to determine optimal startle interstimulus interval

For the first prepulse task the gerbils were asked to detect was a 20-ms gap of silence in an ongoing broadband noise (Fig. 3A). This 20-ms gap prepulse was presented prior to startle stimulus at variable time delays, 5, 10, 20, 40, 80 and 160-ms, in order to determine the ideal time of presentation of prepulse cues relative to the startle stimulus, a value that would be used in the next set of experiments. Detection of the gap was determined by a significant increase in PPI value from a control condition with no gap in the noise (i.e., a 0-ms duration gap) as determined by a one-way ANOVA (see methods). Results revealed a significant effect of gap ISI between the 20-ms gap and the startle stimulus on PPI [F(6,63) = 10.19, p < 0.0001] with post hoc tests showing that gerbils could detect the 20-ms gap at all non-zero ISIs (p’s for all tested ISIs < 0.0007) (Fig. 3B). This same experiment procedure was repeated, but with a 50-ms gap (Fig. 3C). Similarly to the 20-ms gap, the ANOVA revealed that there was also a significant effect of ISI between gap and startle stimulus and PPI [F(6,63) = 12.76, p < 0.0001] with post hoc tests revealing that gerbils could also detect the 50-ms gap at all non-zero ISIs (p’s < 0.0018) (Fig. 3C). Based on these results, we chose to use 80-ms prior to startle as the presentation time for the stimulus in all of the remaining PPI tasks as this value was reliably within the range where PPI plateaued with increasing ISI.

#### 3.2.2 Gap detection in broadband noise

For the next task the ISI between a gap prepulse and the startle stimulus was fixed at 80-ms as determined above, but the gap duration presented in the broadband noise was varied from trial to trial, with values being 1, 3, 5, 10, 15 or 20-ms (Fig. 4A). Detection of the gap was determined by significant increase of PPI value above a control condition where no gap was present in the broadband noise. A one-way ANOVA revealed a significant effect of gap duration on PPI value [F(6,63) = 5.164, p = 0.0002]. Post hoc tests show that gerbils could not detect a gap of 1 (p = 0.8731) or 3-ms (p = 0.1228) but were able to detect gaps of 5 (p = 0.0482), 10 (p = 0.0475), 15 (p = 0.0030) and 20-ms (p = 0.0003) (Fig. 4B). Additionally, the PPI values increased monotonically with increasing gap duration, indicating systematically better detection of longer gaps than shorter gaps similar to a psychometric function that might be measured using operant conditioning methods.

### 3.3 Speaker swap method to measure spatial acuity

#### 3.3.1 Spatial acuity for broadband noise

To first assess spatial acuity, or minimum audible angle (MAA), the broadband noise speaker swap task was used (Fig. 5A). Results from a one-way ANOVA showed a significant effect of speaker swap angle on PPI value [F(5,54) = 11.58, p < 0.0001]. Post hoc tests reveal that gerbils were unable to detect swap angles of 7.5° (p = 0.6851) or 15° (p = 0.4652), but were able to detect swaps of 30° (p = 0.0002), 45° (p < 0.0001) and 90° (p < 0.0001) (Fig. 5B). Additionally, monotonically higher levels of PPI values were produced as a function of angle. Similarly to 0°, compared to an angle of 7.5°, gerbils had significantly better detection of 30° (p = 0.0232), 45° (p = 0.0019) and 90° (p = 0.0015) swap angles. Compared to 15°, gerbils had better detection of angles at 45° (p = 0.0055) and 90° (p = 0.0045). These results show that PPI values systematically and monotonically increased with increasing speaker swap angles, similar to a psychometric function measured using operant conditioning methods. For broadband noise, gerbils exhibited spatial acuity of at least 30°.

#### 3.3.2 Spatial acuity for high- and low-passed noise

Next, gerbils were tested for the ability to utilize high-frequency ILD and low-frequency ITD cues in the spatial acuity speaker swap task (Fig. 6A). A one-way ANOVA run on the high-pass noise speaker swap data shows a significant effect of angle on PPI [F(5,54) = 12.24, p < 0.0001]. Post hoc tests reveal that gerbils were unable to detect speaker swaps of 7.5° (p = 0.9198), but could detect swaps of 15° (p = 0.0024), 30° (p = 0.0037), 45° (p < 0.0001) and 90° (p < 0.0001) (Fig. 6B). Like the broadband noise condition, PPI values, and therefore detection, were larger at wider angles, with PPI values being significantly larger at 15° (p = 0.0417), 45° (p = 0.00143) and 90° (p < 0.0001) degrees compared to 7.5°.

Applied to the low pass results, the one-way ANOVA also showed a significant effect of angle on PPI value [F(5,54) = 2.852, p = 0.0234]. Post-hoc tests showed gerbils were unable to detect speaker swaps of 7.5° (p = 0.2981), 15° (p = 0.7213) or, surprisingly, 90° (p = 0.2115), but could detect swaps of 30° (p = 0.0315) and 45° (p = 0.0279) (Fig. 6C). This significance shows gerbils were able to perform the low pass speaker swap task on the basis of low-frequency ongoing ITDs alone with MAA performance of 30° comparable to that measured in the broadband condition.

#### 3.3.3 Spatial acuity measured within a hemifield

With broadband noise as our swap stimulus, gerbils had similar results when the swap was performed within a hemifield as opposed to across midline. Results from the left hemifield speaker swap showed a significant effect of speaker swap angle on PPI value [F(5,54) = 6.719, p < 0.0001]. Gerbils could detect speaker swaps of 30° (p = 0.0052), 45° (p = 0.0034) and 90° (p = 0.0001), and couldn’t detect angle swaps of 7.5° (p = 0.6833) or 15° (p = 0.4153) (Fig. 7B).

**Fig. 7.**
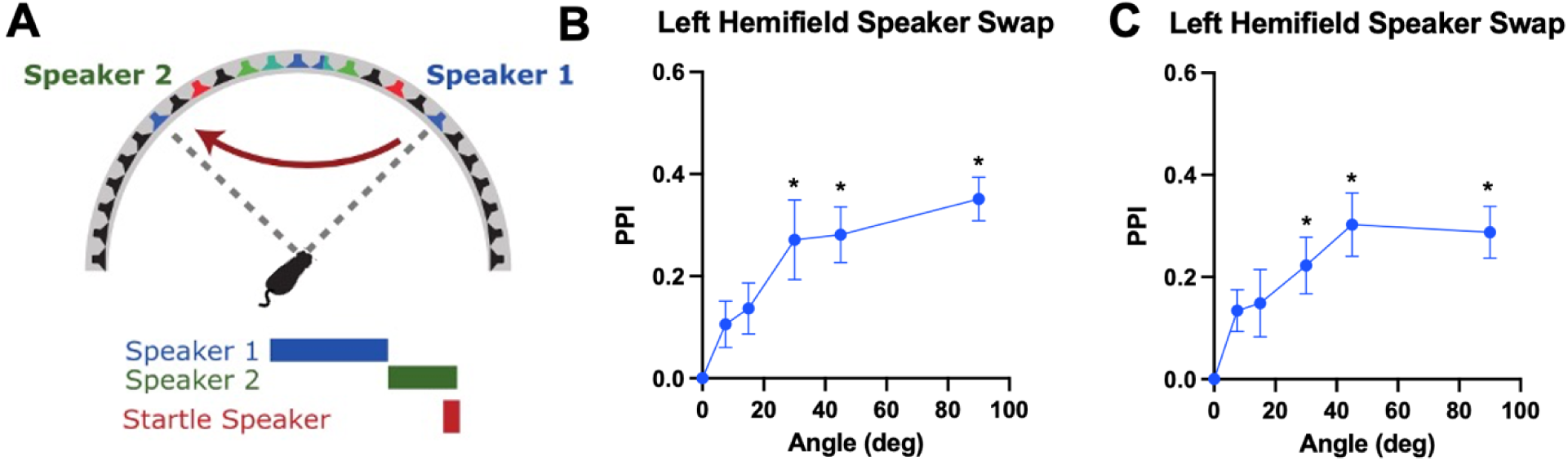
Gerbils can utilize PPI in response to changes in sound source location of broadband noise within a hemifield. **A:** Diagram of the hemifield speaker swap experiment setup. This setup is identical to the broadband noise speaker swap setup, but with animals rotated ±45°, so that each broadband noise speaker swap was contained within either the left or right hemifield of the animal. The angle of the speaker swap was varied. **B:** PPI in response to different speaker swap angles within the left hemifield (7.5, 15, 30, 45 or 90°). Asterisks indicate significant PPI above zero as determined by a one-way ANOVA (*p<0.05). Speaker swap angles ≥ 30° could be detected by gerbils. **C:** PPI in response to different speaker swap angles within the right hemifield (7.5, 15, 30, 45 or 90°). Asterisks indicate significant PPI above zero as determined by a one-way ANOVA (*p<0.05). Speaker swap angles ≥ 30° could be detected by gerbils.

Similar to midline swaps, wider angles had higher levels of detection, as reflected by higher levels of PPI. Gerbils had higher levels of PPI at 90° compared to 7.5° (p = 0.0148) and 15° (p = 0.0462).

To test whether there was any bias towards left versus right hemifield or whether the testing chamber introduced any asymmetric bias in performance due to the different animal orientation, we repeated these experiments, but with the animal rotated to the -45° speaker relative to midline, to test the right hemifield of the animals. The one-way ANOVA on this data showed a significant effect of speaker swap angle on PPI value [F(5,54) = 4.934, p = 0.0009]. Like the left hemifield, post hoc tests revealed that there was no detection at 7.5° (p = 0.4327) or 15° (p = 0.3216), but gerbils could detect speaker swaps of 30° (p = 0.0354), 45° (p = 0.0013) and 90° (p = 0.0026) (Fig. 7C). Thus, gerbils can detect changes in spatial location of sound within a hemifield of at least 30°, comparable to that at the midline, and that our testing chamber did not introduce any left versus right hemisphere bias.

### 3.4 Spatial release from masking task

The final task was meant to mimic a spatial speech in noise task, and tested the ability of gerbils to detect a signal that was spatially separated from a masker. Results from a two-way ANOVA showed that the dB level of the signal [F(2,6) = 88.56, p < 0.0001] had a significant effect on PPI value. Post hoc tests reveal significant differences in PPI across the different signal attenuation levels. PPI in response to the 56 dB signal was significantly higher, and thus more readily detected by the gerbils, than the 44 dB signal at all angles of separation (p < 0.0001 for each angle). PPI in response to the 56 dB signal was also significantly higher, and thus more readily detectable, than the 50 dB signal in all angles (37.5°: p < 0.0001, 22.5°: p = 0.002, 7.5°: p < 0.0001). PPI in response to the 50 dB signal was significantly higher than that elicit in response to the 44 dB signal, but only for maskers presented at 37.5° (p = 0.0069), and 22.5° (p = 0.0018), not at 7.5° (p = 0.7081) (Fig. 8B).

**Fig. 8.**
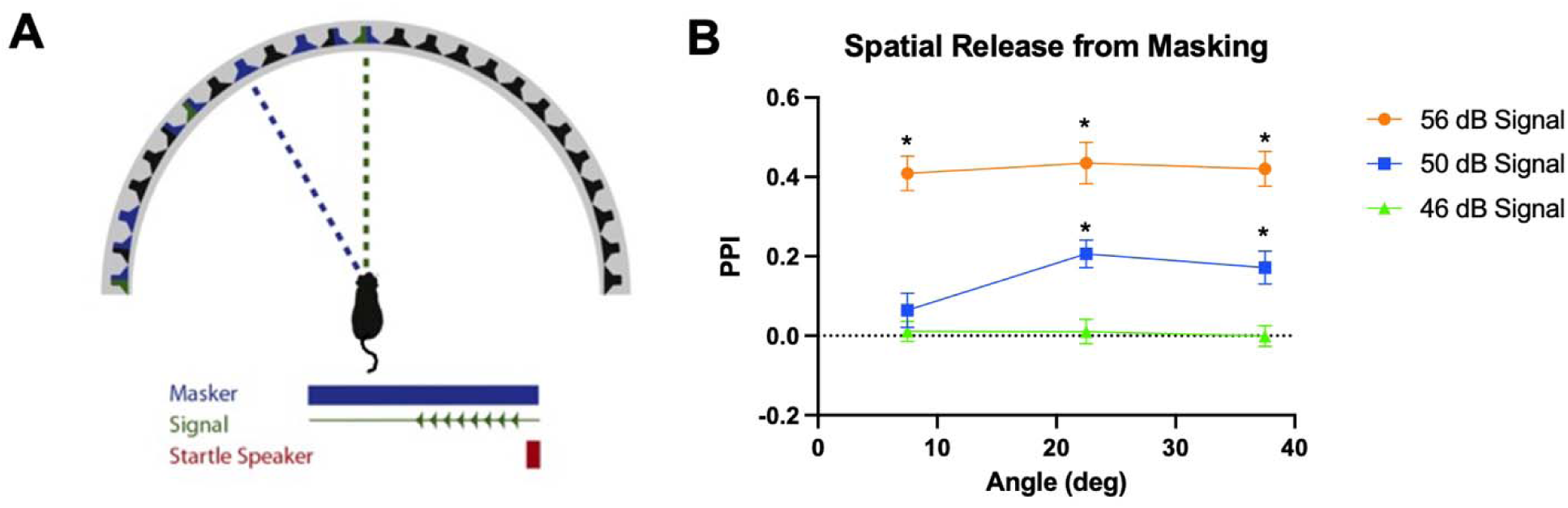
Gerbils can utilize PPI in response to a target presented in spatially separated masking noise. **A:** Diagram of the spatial release from masking experiment setup. Gerbils were held at midline, and a signal consisting of a train of broadband chirps was presented directly ahead of the gerbil. A 70 dB broadband masking noise was also presented along with the chirps. Conditions varied in both the spatial location of the broadband masking noise in relation to the signal, and the attenuation of the signal compared to the broadband masking noise. **B:** PPI in response to different signal levels (44, 50 or 56 dB SPL) and masker spatial locations (7.5, 22.5 or 37.5°). Asterisks indicate significant PPI above zero as determined by a one-way ANOVA for each level tested (*p<0.05). Gerbils could not detect any 44 dB SPL signal, and could detect all 56 dB SPL signals regardless of position relative to masker. At 50 dB SPL, gerbils could detect the signal when the masker was presented 37.5° or 22.5° away from signal, but not when the masker was 7.5°.

We then examined each signal level to see if changes in PPI are elicited from spatial location of the masker compared to the signal through separate one-way ANOVAs. In response to the 56 dB signal, gerbils were able to separate the signal from the masker at all locations presented [F(3,36) = 27.24, p < 0.0001], post hoc tests all significant (p < 0.0001 for each angle). However, there was no difference in the magnitude of PPI between any of the locations tested (p > 0.05) (Fig. 8B). In response to the 44 dB signal level, gerbils were unable to detect the signal against the masker at any location tested [F(3,36) = 0.07623, p = 0.9724], post hoc tests all not significant (p > 0.05 for all angles). Similarly, there was no difference in the magnitude of PPI between any locations tested (p > 0.05) (Fig. 8B).

At 50 dB signal, PPI varied as a function of the location of masker in relation to signal [F(3,36) = 7.714, p = 0.0004]. Post hoc tests reveal that gerbils were able to detect the signal when the masker was presented further away from the signal (37.5°: p = 0.0060, 22.5°: p = 0.0008), but not when the masker was presented closer (7.5°: p = 0.5539). Additionally, we found that PPI was higher in response to masker locations further away from the signal at 22.5° compared to those presented closer to the signal at 7.5° (p = 0.0290) (Fig. 8B). Taken together, the results demonstrate a spatial release from masking in gerbils, particularly for the 50 dB signal.

## 4. Discussion

The gerbil is a well-established model system for the study of neural and behavioral mechanisms of binaural and spatial hearing. Previous groups have shown that gerbils are capable of utilizing PPI for detection of stimuli (Gaese et al., 2009) (Steube et al., 2016). Their work showed that gerbil startle amplitude increased with startle speaker SPL, data which we replicated in our experiments (Fig. 2). We also found an increase in startle amplitude in a 70 dB background noise condition compared to a quiet, 0 dB background condition (Fig. 2), consistent with previous findings in rat startle responses (Cory & Ison, 1979).

Here we used PPI to assess the temporal processing and binaural hearing capabilities of gerbils, based on methods previously used in guinea pigs and mice in our laboratories (Greene et al., 2018) (McCullagh et al., 2020) (Benson et al., 2025). This behavioral test requires no training but rather relies on the fact that mammals, including humans, startle when a loud sound is presented unexpectedly (Fig. 1), and the amount of startle can be quantitatively measured (Fig. 1B). The PPI method leverages the fact that when some aspect of the acoustic sensory environment changes and which is detectable by the animal just before the startle sound is presented, a midbrain reflex circuit (partially) inhibits the startle in a graded way that scales with detectability of the change in the sound (see (Koch, 1999) for review of startle neurobiology).

We made use of this reflex by introducing gaps into ongoing background noise (Fig. 3 and 4), changing the location from which test sounds were emitted (Figs. 5-7), and by presenting a target sound in background noise (Fig. 8).

### 4.1 Gap detection thresholds measured using PPI

In order for PPI to be a useful method, the results produced by it should be comparable to results produced by traditional operant conditioning methods. As demonstrated in Figs 3 and 4, gerbils measured with PPI exhibit gap detection thresholds of ∼3-5 ms, comparable to the ∼2-5 ms thresholds reported using operant conditioning methods (Wagner et al., 2003) (Hamann et al., 2004) (Gleich & Strutz, 2011). These results confirm that our PPI apparatus can measure gap detection thresholds in gerbils consistent with values reported in the literature.

### 4.2 Spatial acuity (MAA) thresholds measured using PPI

Several prior studies have used operant conditioning paradigms to measure gerbil spatial acuity ability (Carney et al., 2011) (Lingner et al., 2012) (Maier & Klump, 2006) (Tolnai et al., 2017) (Lesica et al., 2010) (R. S. Heffner & Heffner, 1988b). For broadband noise sources, these studies have shown spatial acuity ranging from ∼25-38°, which overlaps with the ∼30° measured here using PPI (Fig. 5B).

(Maier & Klump, 2006) reported MAAs with low-pass noise averaging 28.03±5.8° (n = 11 animals across studies), while our measurements (Fig. 6C) indicated thresholds of ∼30°. (Maier & Klump, 2006) also found MAAs of 28.03±5.8° (n = 8) with narrowband noise at 4 and 8 kHz, however this is much higher than the ∼15° found here with high-pass noise at 4 kHz. It is likely that the increased bandwidth in our noise compared to the narrowband noise used by (Maier & Klump, 2006) resulted in additional cues that improved the MAA, such as broader band ILDs, envelope ITDs and spectral shape cues that were not all available to the gerbils with narrowband noise.

While we demonstrate here that PPI methods can be used to measure temporal and spatial hearing abilities in gerbil that are comparable to that measured using operant conditioning methods, some work has suggested that PPI does not allow for the same sensitivity of spatial detection as operant conditioning. For example, (Behrens & Klump, 2016) suggest that PPI in mice may not provide as robust an estimate of MAAs compared to operant conditioning methods, typically underestimating their capabilities. However, here we did not measure MAAs that were larger (i.e., worse spatial acuity) than that reported in the literature for gerbils where operant conditioning methods were used for any of our stimulus conditions, broadband, low- or high-pass noise.

Overall, our results using the PPI method demonstrate spatial acuity in gerbils for broadband as well as low- and high-frequency stimuli that is directly comparable to performance found using traditional operant condition methods.

### 4.3 Sound source localization within a hemifield

It is well known that human listeners can discriminate changes in sound source location not only across the midline but also if the sources are located within a hemifield (i.e. on one side of the head (Mills, 1958)). In humans, MAAs increase (i.e., performance worsens) when discriminating sources away from the midline; however, the decrease in performance can be predicted on the basis of the smaller rate of change with source location of the underlying cues, such as ITDs and/or ILDs, as the reference source for the task moves farther away from the midline (Mills, 1960) (A. D. Brown et al., 2018). The ability to discriminate sound source location changes within a hemifield has been shown in various other species including ferrets (Parsons et al., 1999), the domestic cat (R. S. Heffner & Heffner, 1988a), guinea pigs (Greene et al., 2018) and macaques (C. H. Brown et al., 1982), but not in rats (Kavanagh & Kelly, 1986). Our data demonstrates that gerbils are able to discriminate broadband sound within a hemifield of ∼30° (Fig. 7). These results are consistent with studies of neurons in the auditory cortex, which also indicate an ability to discriminate sound source location within a hemifield in both cats (Stecker et al., 2005) and gerbils (Van Den Wildenberg & Bremen, 2024), but not in rats (Yao et al., 2013).

Thus is appears that the inability of rats to discriminate sound sources within a hemifield is the result of their auditory nervous system, and not because they are poor at spatial acuity per se. In other respects rats have superior abilities to gerbils: rats have MAAs for noise across the midline that are approximately twice as small (i.e., better acuity) as that in the gerbil, ∼15° vs 30°, respectively (H. E. Heffner & Heffner, 1985). The better spatial acuity of the rat relative to the gerbil is due to the fact that the rat’s head is approximately twice as large as that of the gerbil and thus the acoustical cues available for the spatial acuity task, ITDs and ILDs, are much larger for a given change in spatial location (Koka et al., 2008).

Other rodents commonly used for hearing studies include the long-tailed chinchilla and the guinea pig. The chinchilla has a head size similar to that of the rat and thus produces similar location cues (Koka et al., 2011) and similar MAAs of ∼15° for broadband noise, while guinea pigs have head size ∼30% larger than rat and chinchilla (Greene et al., 2014) and therefore better MAAs, ranging from 7.5-15° for noise. Thus gerbils, chinchillas, and guinea pigs are likely to be a superior model to study the neural mechanisms of binaural hearing and sound localization in humans than rats.

### 4.4 Spatial release from masking

The capability of an animal to detect a target within a spatially separated masker condition can be assessed with a spatial release from masking task. While we did not attempt an in-depth investigation of the capabilities of gerbils, our results do demonstrate that when the level of the target was increased and/or the spatial separation between target and masker was increased the signal became more detectable, as indexed by increased PPI (Fig. 8B). When the masker was too intense the gerbils could not detect the target, but when the masker was too faint, the gerbils could always detect the target regardless of the spatial position of the masker. For intermediate masker levels, the detectability of the target depended significantly on the spatial separation of the target and masker, which is a hallmark of spatial release from masking. Thus, out findings support that PPI can be used to measure spatial release from masking in gerbils. While there are no results in the literature in gerbils that are directly comparable to the conditions measured here, there are studies that have shown that gerbils can utilize binaural cues, ITDs and ILDs, to spatially segregate target signals from maskers (Eipert & Klump, 2020).

### 4.5 General discussion and limitations

A potential limitation in our study comes from the physical limitations of our testing apparatus. The animal restraint allowed some side-to-side head movement; however, on average, the animals head was roughly oriented in the appropriate direction since the animal was not able to turn around in the restraint. The large number of repeated testing conditions also allowed us to average together any potential slight variation of head position within the animals. Additionally, we are limited by our speaker positions – we are unable to test angular changes smaller than 7.5° due to the spatial separation of our loudspeakers. Previous work from the lab has navigated this limitation by moving the animal further from the speaker array to create smaller angles (Greene et al., 2018). However, with our gerbils showing MAA thresholds between 15° and 30°, testing for PPI in response to angles <7.5° was deemed unnecessary.

One advantage of the PPI method over operant conditioning is that PPI does not require any training. Operant conditioning experiments can potentially require long periods of training prior to collecting data, followed by many weeks of repeated experimentation to obtain reliable data. Moreover, training on one task, such as spatial acuity, is not generalizable to performance on a different task, such as detecting a signal in spatially separated noise, or tasks of temporal processing, such as gap detection. This makes it near impossible to use operant condition to assess multiple tasks in the same individual. Because of the lack of generalization, operant conditioning is not likely to be as useful for in-depth studies of hearing pathologies resulting from aging (Laumen et al., 2016), noise exposure (Benson et al., 2025), neurodegenerative diseases (e.g. Alzheimer’s (Kurylo et al., 1993)), developmental disorders (e.g. Fragile X (McCullagh et al., 2020), etc., or extensive tests of interventions of therapeutics that may be developed to treat these pathologies. The ability to test groups of animals, such as young vs. older or those noise-exposed or not, treated or not, on several different types of behaviorally-relevant tasks is significant, as human studies have shown that several kinds of listening difficulties related to the conditions listed above arise not only with binaural and spatial hearing (e.g., difficulty understanding speech in noise) but also with mechanisms of general temporal processing. We demonstrated here in *eight* different tests - three gap detection, four MMA and one spatial release from masking - that PPI of the acoustic startle is a robust behavior metric for comprehensive, high-throughput studies of temporal processing and binaural hearing and sound localization, and that PPI can produce results comparable to that measured using operant conditioning methods.

## Conclusion

Here we utilized PPI of the acoustic startle method to rapidly assess the temporal processing as well as binaural and spatial hearing abilities of the Mongolian gerbil. Our results demonstrate that PPI produces a reliable estimate of the behavioral performance of gerbils on gap detection, spatial acuity and spatial release from masking that is directly comparable to performance reported in the literature measured using operant conditioning methods. These results permit future use of PPI methods in gerbils as rapid, high-throughput behavior method to study temporal processing and binaural and spatial hearing dysfunction that exist in many common hearing pathologies and conditions.

## Acknowledgments

We would like to acknowledge Dr. John Peacock for his help with the analysis pipeline. This work was supported by NIH grants F31-DC021622 and R01-DC017924.

## References

1. Allen, P. D., & Ison, J. R. (2010). Sensitivity of the mouse to changes in azimuthal sound location: Angular separation, spectral composition, and sound level. Behavioral Neuroscience, 124(2), 265–277. 10.1037/a0018913

2. Behrens, D., & Klump, G. M. (2016). Comparison of mouse minimum audible angle determined in prepulse inhibition and operant conditioning procedures. Hearing Research, 333, 167–178. 10.1016/j.heares.2016.01.011

3. Benson, M. A., Peacock, J., Sergison, M. D., Stich, D., & Tollin, D. J. (2025). Neural and behavioral binaural hearing impairment and its recovery following moderate noise exposure. Hearing Research, 456, 109166. 10.1016/j.heares.2024.109166

4. Blanca, M., Alarcón, R., Arnau, J., Bono, R., & Bendayan, R. (2017). Non-normal data: Is ANOVA still a valid option? Psicothema, 4(29), 552–557. 10.7334/psicothema2016.383

5. Brown, A. D., Benichoux, V., Jones, H. G., Anbuhl, K. L., & Tollin, D. J. (2018). Spatial variation in signal and sensory precision both constrain auditory acuity at high frequencies. Hearing Research, 370, 65–73. 10.1016/j.heares.2018.10.002

6. Brown, C. H., Schessler, T., Moody, D., & Stebbins, W. (1982). Vertical and horizontal sound localization in primates. Journal of the Acoustical Society of America, 72(6), 1804–1811.

7. Carney, L. H., Sarkar, S., Abrams, K. S., & Idrobo, F. (2011). Sound-localization ability of the Mongolian gerbil (Meriones unguiculatus) in a task with a simplified response map. Hearing Research, 275(1–2), 89–95. 10.1016/j.heares.2010.12.006

8. Cory, R. N., & Ison, J. R. (1979). Persistent effect of noise on the acoustic startle reflex in the rat. Animal Learning & Behavior, 7(3), 367–371. 10.3758/BF03209686

9. Eipert, L., & Klump, G. M. (2020). Interaction of spatial source separation, fundamental frequency and vowel paradigm in Mongolian gerbils. Behavioral Neuroscience, 134(2), 119–132. 10.1037/bne0000356

10. Gaese, B. H., Nowotny, M., & Pilz, P. K. D. (2009). Acoustic startle and prepulse inhibition in the Mongolian gerbil. Physiology & Behavior, 98(4), 460–466. 10.1016/j.physbeh.2009.07.014

11. Gleich, O., & Strutz, J. (2011). The Effect of Gabapentin on Gap Detection and Forward Masking in Young and Old Gerbils. Ear & Hearing, 32(6), 741–749. 10.1097/AUD.0b013e318222289f

12. Goldberg, J. M., & Brown, P. B. (1969). Response of binaural neurons of dog superior olivary complex to dichotic tonal stimuli: Some physiological mechanisms of sound localization. Journal of Neurophysiology, 32(4), 613–636. 10.1152/jn.1969.32.4.613

13. Greene, N. T., Anbuhl, K. L., Ferber, A. T., DeGuzman, M., Allen, P. D., & Tollin, D. J. (2018). Spatial hearing ability of the pigmented Guinea pig (Cavia porcellus): Minimum audible angle and spatial release from masking in azimuth. Hearing Research, 365, 62–76. 10.1016/j.heares.2018.04.011

14. Greene, N. T., Anbuhl, K. L., Williams, W., & Tollin, D. J. (2014). The acoustical cues to sound location in the guinea pig (Cavia porcellus). Hearing Research, 316, 1–15. 10.1016/j.heares.2014.07.004

15. Grothe, B., Pecka, M., & McAlpine, D. (2010). Mechanisms of Sound Localization in Mammals. Physiological Reviews, 90(3), 983–1012. 10.1152/physrev.00026.2009

16. Hamann, I., Gleich, O., Klump, G. M., Kittel, M. C., & Strutz, J. (2004). Age-Dependent Changes of Gap Detection in the Mongolian Gerbil ( Meriones unguiculatus ). JARO - Journal of the Association for Research in Otolaryngology, 5(1), 49–57. 10.1007/s10162-003-3041-2

17. Heffner, H. E., & Heffner, R. S. (1985). Sound localization in wild Norway rats (Rattus norvegicus). Hearing Research, 19(2), 151–155.

18. Heffner, R. S., & Heffner, H. E. (1988a). Sound localization acuity in the cat: Effect of azimuth, signal duration, and test procedure. Hearing Research, 36(2–3), 221–232. 10.1016/0378-5955(88)90064-0

19. Heffner, R. S., & Heffner, H. E. (1988b). Sound localization and use of binaural cues by the gerbil (Meriones unguiculatus). Behavioral Neuroscience, 102(3), 422–428.

20. Joris, P. X., & Yin, T. C. (1995). Envelope coding in the lateral superior olive. I. Sensitivity to interaural time differences. Journal of Neurophysiology, 73(3), 1043–1062. 10.1152/jn.1995.73.3.1043

21. Kavanagh, G. L., & Kelly, J. B. (1986). Midline and lateral field sound localization in the albino rat (Rattus norvegicus). Behavioral Neuroscience, 100(2), 200–205.

22. Koch, M. (1999). The neurobiology of startle. Progress in Neurobiology, 59(2), 107–128. 10.1016/S0301-0082(98)00098-7

23. Koka, K., Jones, H. G., Thornton, J. L., Lupo, J. E., & Tollin, D. J. (2011). Sound pressure transformations by the head and pinnae of the adult Chinchilla (Chinchilla lanigera). Hearing Research, 272(1–2), 135–147. 10.1016/j.heares.2010.10.007

24. Koka, K., Read, H. L., & Tollin, D. J. (2008). The acoustical cues to sound location in the rat: Measurements of directional transfer functions. The Journal of the Acoustical Society of America, 123(6), 4297–4309. 10.1121/1.2916587

25. Kurylo, D. D., Corkin, S., Allard, T., Zatorre, R. J., & Growdon, J. H. (1993). Auditory function in Alzheimer’s disease. Neurology, 43(10), 1893–1893. 10.1212/WNL.43.10.1893

26. Laumen, G., Tollin, D. J., Beutelmann, R., & Klump, G. M. (2016). Aging effects on the binaural interaction component of the auditory brainstem response in the Mongolian gerbil: Effects of interaural time and level differences. Hearing Research, 337, 46–58. 10.1016/j.heares.2016.04.009

27. Lesica, N. A., Lingner, A., & Grothe, B. (2010). Population Coding of Interaural Time Differences in Gerbils and Barn Owls. The Journal of Neuroscience, 30(35), 11696– 11702. 10.1523/JNEUROSCI.0846-10.2010

28. Lingner, A., Wiegrebe, L., & Grothe, B. (2012). Sound Localization in Noise by Gerbils and Humans. Journal of the Association for Research in Otolaryngology, 13(2), 237–248. 10.1007/s10162-011-0301-4

29. Maier, J. K., Kindermann, T., Grothe, B., & Klump, G. M. (2008). Effects of omni-directional noise-exposure during hearing onset and age on auditory spatial resolution in the Mongolian gerbil (Meriones unguiculatus)—A behavioral approach. Brain Research, 1220, 47–57. 10.1016/j.brainres.2008.01.083

30. Maier, J. K., & Klump, G. M. (2006). Resolution in azimuth sound localization in the Mongolian gerbil (Meriones unguiculatus). The Journal of the Acoustical Society of America, 119(2), 1029. 10.1121/1.2159429

31. Maki, K., & Furukawa, S. (2005). Acoustical cues for sound localization by the Mongolian gerbil, *Meriones unguiculatus*. The Journal of the Acoustical Society of America, 118(2), 872–886. 10.1121/1.1944647

32. McCullagh, E. A., Poleg, S., Greene, N. T., Huntsman, M. M., Tollin, D. J., & Klug, A. (2020). Characterization of Auditory and Binaural Spatial Hearing in a Fragile X Syndrome Mouse Model. Eneuro, 7(1), ENEURO.0300-19.2019. 10.1523/ENEURO.0300-19.2019

33. Mills, A. W. (1958). On the Minimum Audible Angle. Journal of the Acoustical Society of America, 30(4), 237–246.

34. Mills, A. W. (1960). Lateralization of High-Frequency Tones. The Journal of the Acoustical Society of America, 32(1), 132–134. 10.1121/1.1907864

35. Owrutsky, Z. L., Benichoux, V., & Tollin, D. J. (2021). Binaural Hearing by the Mammalian Auditory Brainstem: Joint Coding of Interaural Level and Time Differences by the Lateral Superior Olive. In R. Y. Litovsky, M. J. Goupell, R. R. Fay, & A. N. Popper (Eds.), Binaural Hearing (Vol. 73, pp. 113–144). Springer International Publishing. 10.1007/978-3-030-57100-9_5

36. Parsons, C. H., Lanyon, R. G., Schnupp, J. W. H., & King, A. J. (1999). Effects of Altering Spectral Cues in Infancy on Horizontal and Vertical Sound Localization by Adult Ferrets. Journal of Neurophysiology, 82(5), 2294–2309. 10.1152/jn.1999.82.5.2294

37. Pecka, M., Siveke, I., Grothe, B., & Lesica, N. A. (2010). Enhancement of ITD Coding Within the Initial Stages of the Auditory Pathway. Journal of Neurophysiology, 103(1), 38–46. 10.1152/jn.00628.2009

38. Ryan, A. (1976). Hearing sensitivity of the mongolian gerbil, Meriones unguiculatis. Journal of the Acoustical Society of America, 59(5), 1222–1226.

39. Stecker, G. C., Harrington, I. A., & Middlebrooks, J. C. (2005). Location Coding by Opponent Neural Populations in the Auditory Cortex. PLoS Biology, 3(3), e78. 10.1371/journal.pbio.0030078

40. Steenken, F., Heeringa, A. N., Beutelmann, R., Zhang, L., Bovee, S., Klump, G. M., & Köppl, C. (2021). Age-related decline in cochlear ribbon synapses and its relation to different metrics of auditory-nerve activity. Neurobiology of Aging, 108, 133–145. 10.1016/j.neurobiolaging.2021.08.019

41. Steube, N., Nowotny, M., Pilz, P. K. D., & Gaese, B. H. (2016). Dependence of the Startle Response on Temporal and Spectral Characteristics of Acoustic Modulatory Influences in Rats and Gerbils. Frontiers in Behavioral Neuroscience, 10. 10.3389/fnbeh.2016.00133

42. Tollin, D. J. (2003). The Lateral Superior Olive: A Functional Role in Sound Source Localization. The Neuroscientist, 9(2), 127–143. 10.1177/1073858403252228

43. Tolnai, S., Beutelmann, R., & Klump, G. M. (2017). Exploring binaural hearing in gerbils (Meriones unguiculatus) using virtual headphones. PLOS ONE, 12(4), e0175142. 10.1371/journal.pone.0175142

44. Turner, J. G., Parrish, J. L., Hughes, L. F., Toth, L. A., & Caspary, D. M. (2005). Hearing in Laboratory Animals: Strain Differences and Nonauditory Effects of Noise. Comparative Medicine, 55(1), 12.

45. Van Den Wildenberg, M. F., & Bremen, P. (2024). Heterogeneous spatial tuning in the auditory pathway of the Mongolian Gerbil ( *MERIONES UNGUICULATUS* ). European Journal of Neuroscience, 60(5), 4954–4981. 10.1111/ejn.16472

46. Wagner, E., Klump, G. M., & Hamann, I. (2003). Gap detection in Mongolian gerbils (Meriones unguiculatus). Hearing Research, 176(1–2), 11–16. 10.1016/S0378-5955(02)00643-3

47. Yao, J. D., Bremen, P., & Middlebrooks, J. C. (2013). Rat primary auditory cortex is tuned exclusively to the contralateral hemifield. Journal of Neurophysiology, 110(9), 2140– 2151. 10.1152/jn.00219.2013

48. Yin, T. C. T., & Chan, J. C. (1990). Interaural time sensitivity in medial superior olive of cat. Journal of Neurophysiology, 64(2), 465–488.

49. Yin, T. C. T., Smith, P. H., & Joris, P. X. (2019). Neural Mechanisms of Binaural Processing in the Auditory Brainstem. In R. Terjung (Ed.), Comprehensive Physiology (1st ed., pp. 1503–1575). Wiley. 10.1002/cphy.c180036

